# TDP-43 pathology is linked to motor neuron loss and is independent of stress granules *in vivo*

**DOI:** 10.64898/2026.02.11.705439

**Authors:** Alicia Dubinski, Ala Ferdi, Mariam Choughari, Holly Spence, Arpita Adhikary, Carolane Fauchon, Melissa Touati, Myriam Gagné, Martha Liu, Sarah Peyrard, Jenna Gregory, Christine Vande Velde

## Abstract

Nuclear depletion and cytoplasmic aggregation of TDP-43 define a pathological signature across amyotrophic lateral sclerosis (ALS), frontotemporal dementia (FTD), Alzheimer’s disease, and limbic-predominant age-related TDP-43 encephalopathy (LATE). Stress granule persistence and chronic activation of the integrated stress response (ISR) have been proposed to trigger this pathology, yet clinical trials targeting these pathways have failed despite robust target engagement suggesting that the prevailing model may be incomplete. Here, we use a physiologically relevant recurrent hyperthermia paradigm to directly test the relationship between stress granules and TDP-43 pathology *in vivo*. We find that RNA-binding proteins typically associated with stress granules persist as dynamic, phase-separated cytoplasmic assemblies in spinal motor neurons of both wild-type and mutant TDP-43 mice. These structures resolve spontaneously and are spatially distinct from TDP-43 puncta. Strikingly, in mutant TDP-43 mice with a compromised acute stress granule response, stress exposure provokes TDP-43 nuclear export and cytoplasmic deposition, culminating in selective loss of spinal α-motor neurons after recurrent stress. Our results reveal that TDP-43 nuclear clearance and cytoplasmic aggregation can occur independently of stress granules *in vivo*, overturning a central assumption of TDP-43 pathogenesis. This paradigm shift reframes the mechanistic link between cellular stress and TDP-43 pathology, providing a new perspective for therapeutic strategies related to ISR modulation.

## Background

There is an immediate need in the field to validate whether targeting the stress granule pathway is a valid therapeutic strategy for ALS/FTD and other neurodegenerative diseases. Stress granules form as part of the integrated stress response (ISR) and are proposed to act as triage sites for polyadenylated (polyA) mRNA as well as signaling hubs for pathways such as apoptosis^1,2^. Importantly, they are transient cytoplasmic structures that form via liquid-liquid phase separation during external insults, and disassemble rapidly following abatement of the stress exposure^3^. In ALS, chronic upregulation of the ISR has been observed in both patient samples and animal models^4,5^, and RNA binding proteins (RBPs) classically catalogued as stress granule proteins, such as TIA-1 and HuR, have been reported to co-label cytoplasmic TDP-43 aggregates in post-mortem tissues^6,7^. Due to a plethora of data primarily derived from *in vitro* overexpression systems, a prevailing hypothesis in the field is that persistent stress granules, caused by failed stress granule disassembly, act as seeds that precede TDP-43 aggregates and are neurotoxic^8–10^. Moreover, recent work has suggested that modulating stress granule dynamics with the small molecule lipoamide can reduce the axonal damage observed in mutant FUS expressing iPSC-derived motor neurons as well as improve behavioral phenotypes in mutant TDP-43 and FUS *Drosophila* models^11^. Recent clinical efforts to dampen the ISR using eIF2B agonists (DNL-343 & Fosigotifator), and thus abolish persistent stress granules, have failed to show clinical improvements in ALS patients, despite clear target engagement for DNL-343^12,13^. More recently, the protective nature of stress granules has been investigated. For example, stress granules serve a protective role in fly models of ALS and FTD, as their dysfunction is associated with more severe disease outcomes^14,15^. In addition, in a mouse model expressing near-physiological levels of the ALS-linked TDP-43 mutation M337V, stress granule assembly is defective in motor neurons *in vivo* and is not correlated to TDP-43 translocation events^16^. It remains unknown whether stress granule dysfunction (protective or toxic) is linked to ALS pathogenesis.

It has been proposed that ALS onset is a combination of genetic predisposition, aging, and environmental insults^17^. While many transgenic mouse models that have been generated via TDP-43 overexpression (accumulating protein levels that are >2-fold over the endogenous) show clear motor phenotypes and neurodegeneration, knock-in models tend to show very mild and late-onset behavioral phenotypes, many of which are more closely associated with FTD^18–21^. While the field has been dominated by studies demonstrating a gain of toxicity owing to TDP-43 cytoplasmic aggregation, there has a recent shift towards a TDP-43 loss-of-function (LOF) hypothesis. Specifically, the loss of TDP-43 mediated repression of cryptic exons, such as those found in *STMN2* and *UNC13A*, which correlates positively with TDP-43 pathology in patients^22–26^.

There is a clear need for improved TDP-43 mouse models that can advance our understanding of the relationship between TDP-43 LOF, TDP-43 aggregation, and disease pathogenesis^19,27^. One way to accomplish this is by challenging existing TDP-43 models with an environmental insult (“two hit” models) to induce more severe molecular and behavioral phenotypes. Along this line, a recent study has shown that challenging a mouse model expressing the ALS-associated *C9orf72* repeat expansion with repetitive traumatic brain injury (TBI) leads to TDP-43 translocation, reduced neuron density, and worsened behavioral phenotypes^28^. Developing “two hit” models that incorporate genetic predisposition and environmental insult, rather than models that utilize supraphysiological expression of TDP-43, is anticipated to improve translation of future therapeutics.

In this study, we adapted our previously published acute hyperthermia model to determine if stress granules could persist *in vivo* and whether this associated with TDP-43 pathology or motor neuron death. In addition, we aimed to evaluate how this environmental stress influenced TDP-43 translocation and aggregation in TDP-43^M337V^ mice, which display defective stress granule assembly and increased TDP-43 nuclear clearance during acute hyperthermia. Our data demonstrate that stress granule proteins, but not cytoplasmic demixed polyadenylated mRNA, persist in the motor neurons of mice exposed to whole body recurrent hyperthermia, but this is reversible and not associated with neuronal survival. Our data also indicate that cytoplasmic TDP-43 foci are spatially independent of these structures. Lastly, we show that recurrent heat stress exposures in this TDP-43 mouse model induces TDP-43 nuclear clearance, accumulation of cytoplasmic TDP-43 puncta, loss of TDP-43 function, and selective death of alpha motor neurons.

## Methods

### Animals

The mouse line used was the TDP-43^M337V^ BAC transgenic mouse^29^. Only males were used for all recurrent hyperthermia experiments in the manuscript, as they are reported to have a stronger phenotype in the TDP-43^M337V^ mouse line^29^. For data with non-transgenic mice, all mice were littermates with no transgene expression in the TDP-43^M337V^ colony. Mice used in Figure 1 were a combination of B6/N mice (Charles River) and NTg littermates from TDP-43^M337V^ mice.

**Figure 1.**
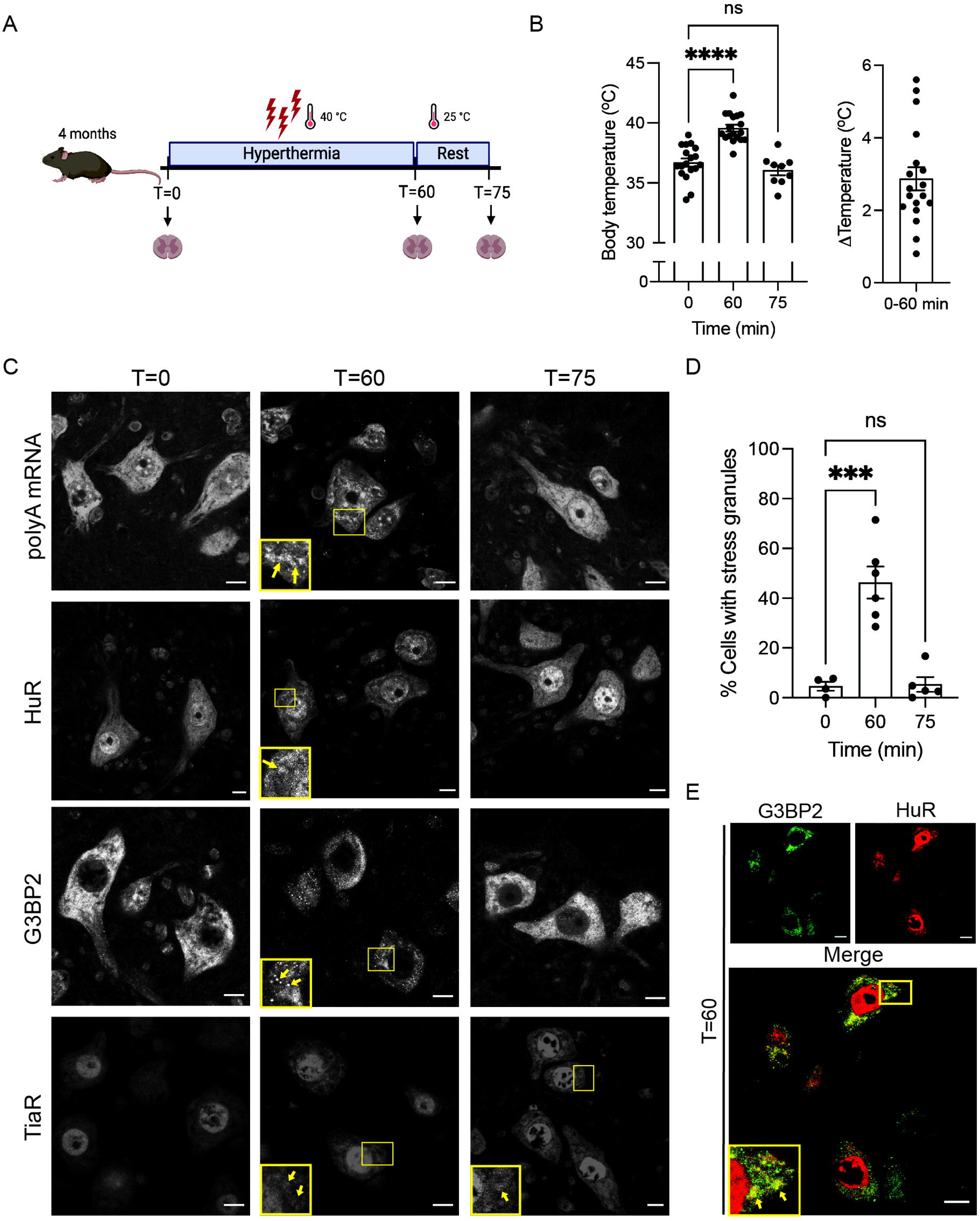
Stress granules assemble and disassemble in vivo. (**A**) Schematic of stress paradigm with awake, restrained mice. Mice were perfused at T=0, T=60, and T=75 for analysis. (**B**) Temperature data for mice at T=0 (N=17), T=60 (N=17), and T=75 (N=9). Combined from NTg and TDP-43^M337V^ genotypes. Absolute temperature at each time point and increase during 60 min stress are presented. (**C**) Representative fluorescence *in situ* hybridization (FISH) images for detection of polyadenylated mRNA in spinal motor neurons, and immunofluorescence (IF) images for stress granule resident proteins HuR, G3BP2, and TiaR. Scale bar 10 µm. Insets show granular signals for stress granule markers. Arrows show persistent TiaR puncta after 15 mins recovery (T=75). **(D)** Quantification of cells with stress granules at different time points using FISH images; T=0 (N=4), T=60 (N=6), T=75 (N=4). **(E)** Representative IF showing overlap of HuR and G3BP2 at 60 mins. Scale bar 10 µm.

### Recurrent hyperthermia treatment

For the 60 mins hyperthermia treatment, awake mice were weighed and starting core temperatures were taken immediately (PhysioSuite, Kent Scientific). Mice were placed in a restraining device (Stoelting) and lowered into their cage. Cages were placed into an incubator set to 40°C supplemented with 1L/min O_2_ and a 60 min timer was started. Mice were observed every 10 mins to ensure they were moving adequately and not in distress. After 60 mins of hyperthermia, mice were removed from restrainers, and the internal body temperature was taken immediately. Mice were then given free access to wet food and water for the 15 min rest period. This was repeated immediately a for a second 60 min stress. After the second 15 min rest period (total of 150 mins), mice were given 1.0 mL of saline for hydration in addition to diet-gel and wet food to prevent weight loss. The 150 min hyperthermia paradigm was repeated twice per week for 4 weeks. The next day, mice were sacrificed by perfusion with 4% PFA or for tissue collection by flash freezing in liquid N_2_. Tissues were subsequently cryopreserved, embedded in OCT, and sectioned. Mice in the recovery paradigm (12M + 1M hyperthermia + 4M recovery) were monitored biweekly throughout the recovery period. Following this, they were sacrificed by perfusion with 4% PFA and processed. Mice were weighed three times per week throughout the hyperthermia and biweekly for the recovery period.

### Immunofluorescence and fluorescence in situ hybridization (FISH)

Floating sections of lumbar spinal cords (L3-L5, 30μm) were used for staining. For all labellings, sections were washed in Sorenson’s PBS (13mM NaH_2_PO_4_, 87mM Na_2_HPO_4_, pH 7.6) then blocked with 3% BSA followed by incubation of primary antibodies (TDP-43 Proteintech 10782-2-AP; 1/200, TDP-43 R&D Systems MAB7778; 1/200, TiaR BD Biosciences 610352; 1/100, HuR SantaCruz sc-5261; 1/300, G3BP2 Cedarlane A302-040A; 1/100, TDP-43 S409/410 Cedarlane 829901; 1/200, p62 Sigma P0067; 1/100), human-specific TDP-43 #60019-2-Ig; 1:10,000) overnight at 4°C. The following day, spinal cords were washed and incubated with fluorescently conjugated secondary antibodies (1/200) to the desired species, stained with Hoescht, and mounted on slides with ProLong Antifade reagent (Invitrogen). For fluorescence in situ hybridization, sections were washed in 5X SSC buffer, acetylated with acetic anhydride (3μL/mL), permeabilized with Triton-X-100 (0.5% in 5X SSC), then incubated in hybridization buffer (0.1M TEA pH8, 25% formamide, 1mg/mL yeast RNA, 0.01% BSA, 10% dextran sulfate) for 2 hr at 37°C. The oligo-dT probe (1.3ng/μL, Texas Red) was diluted in hybridization buffer then added to spinal cords for incubation overnight at 37°C. The following day, spinal cords were diluted and washed in 5X SSC buffer, stained with Hoescht, and mounted with ProLong Antifade reagent (Invitrogen). Images were acquired with a 63X oil (1.7 NA) objective on a confocal microscope (SP5; Leica) using LAS-X software (Leica).

### BaseScope

The Sort1 exon 17B BaseScope probe and assay kit were purchased from ACD Bio and used according to the manufacturer’s instructions (Advanced Cell Diagnostics, USA). Coronal brain sections (8 μm) were mounted onto adhesive slides and fixed in 4% paraformaldehyde (PFA) for 1 h prior to BaseScope processing. Sections were incubated with hydrogen peroxide, subjected to target retrieval for 5 min, and treated with Protease III. Hybridization with the Sort1 exon 17B probe was performed for 2 hr at 40°C, followed by sequential amplification using reagents 1–8. Sections were counterstained with hematoxylin, mounted with VectaMount, and imaged using a Leica brightfield microscope.

### Labelling with TDP-APT

Coronal sections (8 μm) were mounted onto adhesive slides, fixed with 4% PFA for 1 hr and washed in PBS-T (0.05% Tween) for 10 mins. Biotin blocking was carried out using a kit (Abcam, ab64212) as per the manufacturer’s guidelines followed by washing in PBST (0.01% Tween). A protein block was performed using the Novolink Polymer detection system (Leica Biosystems, Newcastle, UK). TDP-43 aptamer (1/100, TDP-43^Apt^ CGGUGUUGCU with a 3’ Biotin-TEG modification, ATDBio, Southampton, UK) prepared in protein block was applied to the tissue and incubated for 3 h at 4 °C followed by incubation with 4% PFA overnight at 4 °C. Sections were incubated with primary and secondary antibodies (1/100 goat-anti biotin Ab6652, 1/200 A647 donkey anti-goat Ab150135) then washed and mounted with Vectashield medium with DAPI (Vector labs, H-1200). Images were acquired with a ZEISS AxioScan Z1 slide scanner at 20X magnification microscope at 20X magnification.

### Motor neuron counting

Motor neuron counting was done using immunofluorescence on lumbar spinal cords (L3-L5). The same immunofluorescence protocol was followed using ChAT (Millipore AB144P; 1/100) and NeuN (Thermo Fisher PA5-78499; 1/500) primary antibodies. Between 7-10 sections, at least 150 μm apart, were imaged per mouse. Alpha motor neurons were stained for both ChAT and NeuN, while gamma motor neurons were stained positive for ChAT but do not have a NeuN signal ^31^. Images were acquired with a 20X (1.25 NA) objective and analyzed using Cell Counter on ImageJ.

### Image Analysis

Neurons were scored as positive for stress granules if they displayed at minimum 2 puncta that measured at least 0.15 μm^2^. A minimum of 15 motor neurons were analyzed per mouse. Cells with TiaR or TDP-43 nuclear depletion were counted manually. Images were taken in lumbar spinal cords focused on the ventral horn, and a cell was counted if it had a nuclear size of at least 20μm^2^. For TiaR puncta, cells were scored as positive if they displayed a minimum of 5 puncta measuring at least 0.15 μm^2^. For TDP-43 puncta, cells were considered positive if they had a minimum of 5 puncta measuring 0.15 μm^2^, or a single punctum that was a minimum of 1 μm^2^, due to the background of the antibody staining. All image analysis was done using ImageJ.

### Cell culture

Mouse neuroblastoma cells (N2A, gift from Nicole Leclerc) were maintained in DMEM at 37C and 5% CO2. For siRNA transfection, non-targeting and TDP-43 targeting siRNAs (Dharmacon #D-001810-10-20, #L-040078-01-0020) were transfected using oligofectamine for 5 hours (Invitrogen 12252-011) and collected for RNA extraction 72 hours later.

### RT-PCR

Total RNA was extracted from cell pellets (N2A) using QIAGEN RNeasy mini kit according to the manufacturer instructions. RNA was isolated from mouse spinal cords using trizol/chloroform followed by isopropanol preceiptation. Residual genomic DNA was removed using the TURBO DNA-*free*™ Kit (Invitrogen #AM1907). RNA concentration was measured and equal amounts of RNA were used to synthesize cDNA with a reverse transcriptase kit (Qiagen). cDNA was amplified using primers targeting Adnp2 cryptic exon (339 bp, F: CTTGACAACATCAGGAAGGTGC, R: TTGTCCGGTATCTCGCTTTCT), Synjbp2 cryptic exon (224 bp, F: CTCCAACGACAGTGGCATCT, R: TCTTCCTGAGGACCTCCGTT), Usp15 cryptic exon (356 bp, F: GGTCCCTCTACTCCTAAGTCCC, R: TGGCTGTT CATTGTTTCTTCCAG), and Sort1-17B (350 bp, F: CAAATGCCAAGGTGGGATGAA, R: TTGAATCCAAAGCCTCTACGCC)^32,33^. PCR was performed using Phusion Plus green Master Mix (Thermo Scientific #F632L) with a modified touchdown PCR protocol. Products were visualized by agarose gel electrophoresis using SYBR safe.

### Statistics

T-tests and ANOVA analysis were used to measure statistical significance (Prism, GraphPad). A p-value lower than 0.05 was considered significant.

## Results

### Stress granule assembly and disassembly in vivo

We have previously shown that mice bearing the TDP-43^M337V^ mutation have a robust defect in stress granule formation^16^. However, as the acute hyperthermia protocol is a terminal procedure^16^, this precluded the evaluation of stress granule disassembly. Thus, we designed a new protocol in which awake, mildly restrained mice were maintained for 60 mins in an incubator set to 40°C to induce stress granule formation in motor neurons (***Fig. 1A***). The internal body temperature of the mice was raised approximately 2.5°C (39°C avg internal temp) after 1 hr and normalized within 15 mins of their return to ambient temperature (***Fig. 1B***). Mice were continuously monitored throughout the 1 hr to ensure adequate respiration and absence of distress. After 1 hr, mice were given free access to food and water for a 15-min recovery period. Premised on our prior work demonstrating that stress granules can be monitored in lumbar spinal cord neurons (L3-L5) by following the partitioning of cytoplasmic polyA mRNA (***Fig. 1C***), which is an obligate component of all stress granules, we established that this recurrent heat stress paradigm induced stress granule formation in ∼50% of neurons at 60 mins, which fully disassembled after 15 mins of recovery (***Fig. 1D***). In addition to polyA mRNA and consistent with our previous work, the stress granule proteins TiaR, G3BP2, and HuR showed cytoplasmic demixing which also resolved after 15 mins of recovery (***Fig. 1C***), with HuR and G3BP2 showing colocalization (***Fig. 1E***). Note, we observed that not all protein markers were equivalent, with cytoplasmic TiaR puncta tending to persist after the 15 mins recovery period, while HuR and G3BP2 completely resolved (***Fig. 1D, arrows***). The variable protein patterns further validates the use of oligo-dT to track canonical stress granules. Overall, this revised heat stress paradigm facilitates the evaluation of both stress granule assembly and disassembly in mouse motor neurons *in vivo*.

### Stress granule proteins, but not polyA mRNA, remain phase separated after recurrent hyperthermia

Consistent with our prior work using acute heat stress^16^, motor neurons in TDP-43^M337V^ mice failed to form stress granules following 60 mins at 40°C (***Supplementary Fig. 1***). Using mice aged 4 months (4M) and 12 months (12M), ages at which we have previously established that stress granule formation in motor neurons is intact *in vivo*^16^, we performed two rounds of 60 mins stress followed by a 15 mins recovery period twice per week for four weeks in TDP-43^M337V^ transgenic mice and their non-transgenic (NTg) littermates (***Fig. 2A***). Over the course of the stress protocol, the average temperature increase per 60 mins session of hyperthermia was 1.8°C and 3.0°C in 4M and 12M animals, respectively (***Supplementary Fig. 2A, B***), and the weight of the mice remained stable (***Supplementary Fig. 2C, D***). After 4 weeks of recurrent heat stress, polyA mRNA granules were not detected in either 4M or 12M old mice of either genotype (***Fig. 2B***), as expected since stress granules are transient structures^35^. However, at both ages and in both genotypes, TiaR puncta were detected following recurrent stress exposure in a subset of lumbar spinal motor neurons (***Fig. 2C, insets***). In 4M NTg mice, TiaR puncta were observed in 66% of neurons in mice exposed to recurrent hyperthermia, in contrast to 37% of neurons in sham-treated animals (***Fig. 2D***). At 12M, NTg mice exposed to recurrent hyperthermia had significantly fewer neurons with TiaR puncta (30%). TDP-43^M337V^ mice largely mirrored NTg mice, with fewer neurons displaying persistent TiaR puncta in 12M recurrently stressed animals compared to 4M mice (***Fig. 2E***). Given that stress granule formation is preserved in NTg, but impaired in TDP-43^M337V^ mice, this suggests that the formation of persistent TiaR puncta is independent of stress granule formation. Additionally, we observed persistent HuR and G3BP2 cytoplasmic puncta in NTg mice (***Supplementary Fig. 3***), suggesting a general persistence of stress granule proteins after recurrent hyperthermia, and that the persistence of TiaR puncta is not specific. TiaR is reported to also translocate to the cytoplasm following a stress exposure *in vitro*^36,37^, thus we also quantified neurons with loss of nuclear TiaR after hyperthermia. Translocation of TiaR from the nucleus to the cytoplasm was significantly higher in stressed 4M NTg mice compared to sham-treated control mice, and was unchanged between conditions in 12M mice **(*Fig. 2F*)**. In contrast, 4M and 12M TDP-43^M337V^ mice displayed a very small proportion of neurons with TiaR nuclear clearance (3-6%) and it was not stress-dependent (***Fig. 2G***). In summary, we report that stress granules, as labeled with polyA mRNA, resolve following recurrent hyperthermia, while cytoplasmic foci containing the RNA binding proteins TiaR, HuR, and G3BP2 do not. Further, we conclude that these persistent immunoreactive structures represent protein assemblies that are inconsistent with features of *bona fide* stress granules.

**Figure 2.**
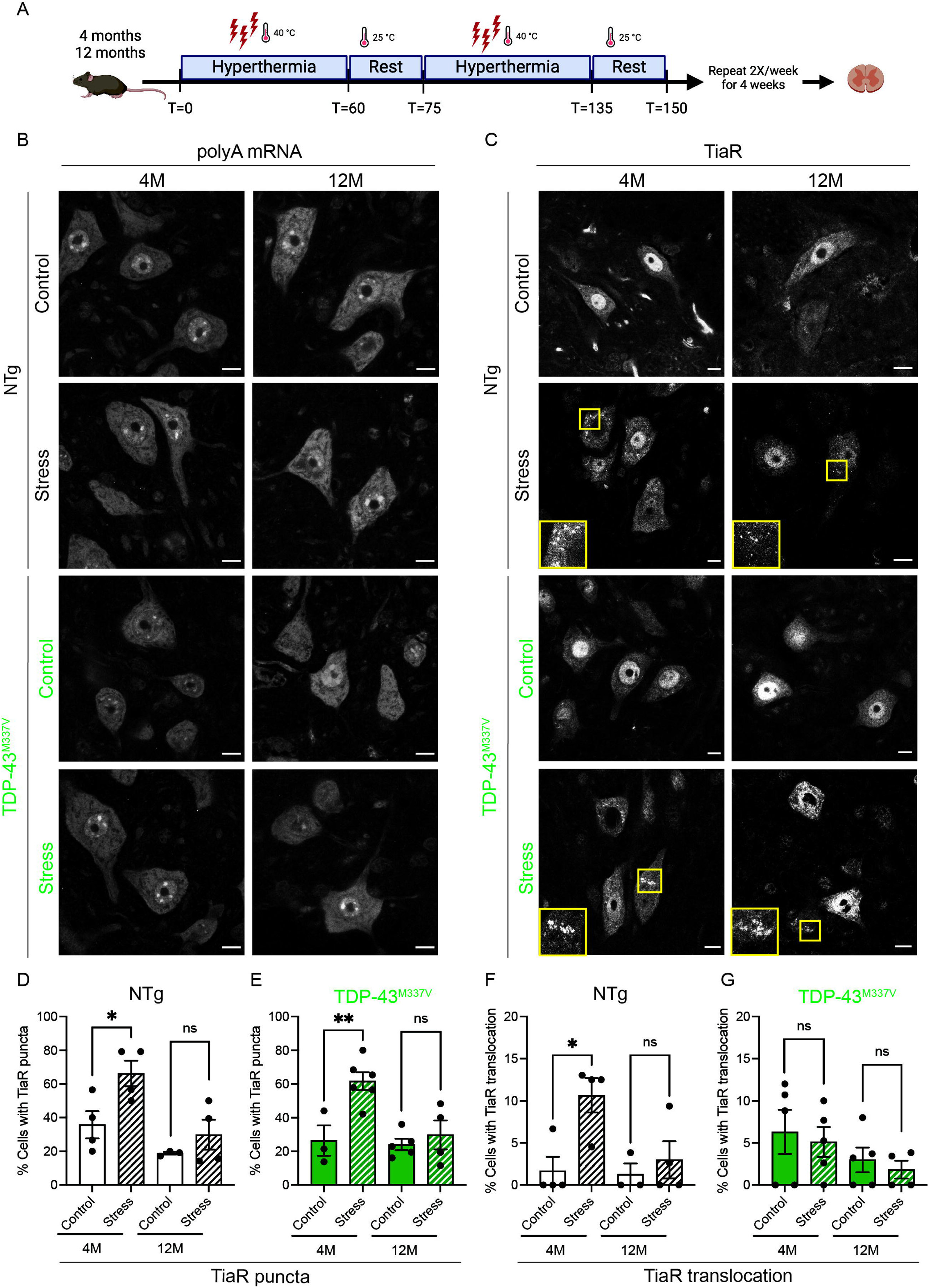
TiaR puncta, but not persistent stress granules, are present after recurrent hyperthermia. **(A)** Schematic of stress paradigm for recurrent hyperthermia starting at 4M and 12M, performed in NTg and homozygous TDP-43^M337V^ mice. **(B)** Representative FISH images for polyA mRNA in spinal motor neurons at 4M and 12M. Scale bar 10 µm. **(C)** Representative TiaR images from spinal motor neurons at 4M and 12M. Scale bar 10 µm. Insets show persistent TiaR puncta in motor neurons after hyperthermia. **(D-E)** Quantification of neurons with TiaR puncta at 4M and 12M after stress; NTg: Control 4M (N=4), Stress 4M (N=4), Control 12M (N=3), Stress 12M (N=4); TDP-43^M337V^: Control 4M (N=3), Stress 4M (N=6), Control 12M (N=5), Stress 12M (N=4). **(F-G)** Quantification of TiaR translocation in neurons at 4M and 12M. Cells were considered to have translocation when the nucleus showed no TiaR signal; NTg: Control 4M (N=4), Stress 4M (N=4), Control 12M (N=3), Stress 12M (N=4); TDP-43^M337V^: Control 4M (N=5), Stress 4M (N=5), Control 12M (N=5), Stress 12M (N=4). One-way ANOVA was used to determine statistical significance.

### TDP-43^M337V^ mice develop neuronal cytoplasmic TDP-43 puncta in response to recurrent stress

It has been suggested that TDP-43 nuclear egress is part of a normal physiological response to stress^38–40^. Similar to what we have previously reported in a model of acute heat stress in this mouse model, TDP-43 nuclear clearance was observed in ∼15% of the lumbar motor neurons of 4M NTg mice exposed to recurrent hyperthermia and was variable in 12M mice (***Fig. 3A, C*)**. This stress-dependent relocalization of TDP-43 was absent in TDP-43^M337V^ mice exposed to recurrent hyperthermia and instead remained similar to sham-treated mice at both ages examined (***Fig. 3B, D***). To evaluate the presence of cytoplasmic TDP-43, we used two different commercial antibodies with reactivity to mouse and human TDP-43 (***Fig. 3E***). Specifically, a monoclonal antibody from R&D Biosystems (MAB7778) displayed larger cytoplasmic inclusions, while a polyclonal antibody available from Proteintech (10782-2-AP) showed several small cytoplasmic puncta within lumbar spinal neurons (***Fig. 3E***). Using the R&D Biosystems antibody for analysis, we detected TDP-43 puncta in ∼35% of lumbar spinal neurons in 12M TDP-43^M337V^ mice exposed to recurrent hyperthermia, while they were detected in only ∼15% of neurons from age-matched NTg mice (***Fig. 3F***). Furthermore, it was evident that TDP-43 nuclear clearance and cytoplasmic puncta were almost completely independent events, with only 1.5% of neurons (6/400) showing co-occurrence of these events (***Fig. 3G***). Analysis of TDP-43 nuclear clearance and TDP-43 cytoplasmic puncta with the Proteintech antibody yielded similar results (***Supplementary Fig. 4A-C***). To further assess TDP-43 pathology, we also labelled the motor cortex of 12M TDP-43^M337V^ mice with an aptamer that recognizes pathological TDP-43^41^. We observed that while both stressed mice and sham-treated controls have TDP-43 pathology, there was increased signal after hyperthermia (***Fig. 3H***). As it is unclear whether the TDP-43 phenotypes we observed are associated with loss of TDP-43 function, we assessed the levels of three TDP-43 dependent cryptic exons in addition to splicing of *Sort1*, using N2A cells treated with TDP-43 siRNA as a positive control. Using RT-PCR of spinal cord lysates, no splicing events that known to be associated with TDP-43 loss of function were observed at baseline in TDP-43^M337V^ mice, nor following recurrent stress in bulk tissues (***Supplementary Fig. 4D***). However, BaseScope analysis of *Sort1 17B* in the cortex of TDP-43^M337V^ mice demonstrated a marked increase after recurrent hyperthermia, consistent with a stress-induced loss of neuronal TDP-43 splicing function (***Fig. 3I***). Taken together, in response to recurrent stress, we observe cytoplasmic TDP-43 puncta in motor neurons, which are exacerbated by stress and genotype. Our data suggests that these changes are associated with TDP-43 loss of function.

**Figure 3.**
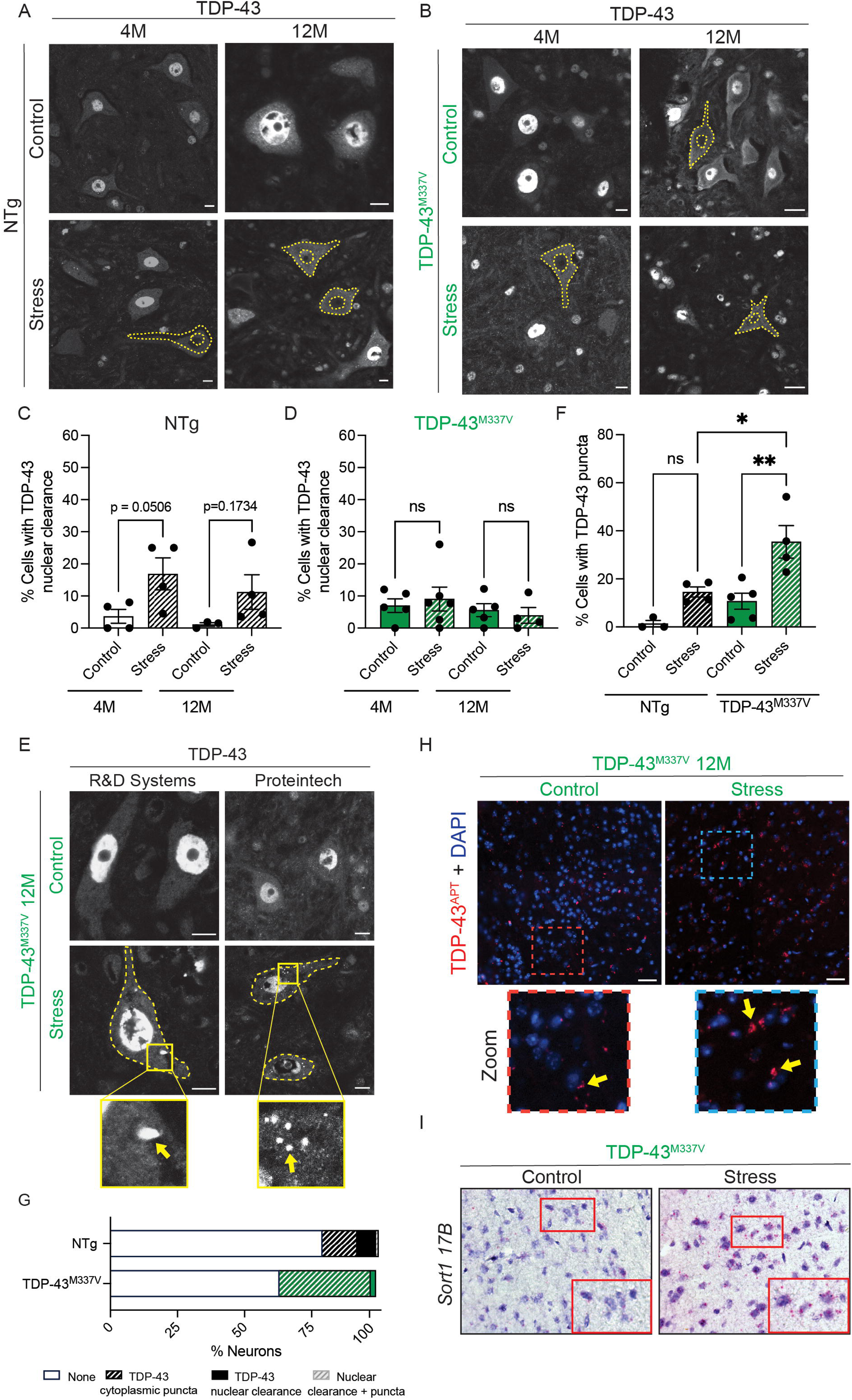
TDP-43 translocation and puncta occur in motor neurons after recurrent hyperthermia. **(A-B)** Representative images of TDP-43 translocation and at 4M and 12M in NTg and TDP-43^M337V^ mice after recurrent stress. Scale bars are 10 µm. Yellow outlines depict neurons with TDP-43 nuclear clearance. **(C-D)** Quantification of cells with TDP-43 nuclear clearance at 4M and 12M. NTg: Control 4M (N=4), Stress 4M (N=4), Control 12M (N=3), Stress 12M (N=4); TDP-43^M337V^: Control 4M (N=5), Stress 4M (N=5), Control 12M (N=5), Stress 12M (N=4). **(E)** Representative IF images TDP-43 cytoplasmic inclusions detected with two different antibodies (R&D Systems #MAB7778, Proteintech #10782-2-AP) after 12M recurrent stress. Yellow arrows point to TDP-43 puncta. Scale bars are 10 µm. **(F)** Quantification of neurons with TDP-43 puncta at 12M using R&D Biosystems antibody. NTg: Control 12M (N=5), Stress 12M (N=5), TDP-43^M337V^: Control 12M (N=5), Stress 12M (N=4). **(G)** Analysis of relationship between TDP-43 nuclear clearance and puncta formation. NTg (N=119); TDP-43^M337V^ (N=111). **(H)** Representative images of labelling with TDP-43^APT^ to detect TDP-43 pathology in the motor cortex. TDP-43^M337V^ 12M Control N=2, Stress N=2. **(I)** Representative BaseScope images with Sort1 17B probe in cortex samples of sham-treated or hyperthermia-treated 12M TDP-43^M337V^ mice.

### Neuronal TDP-43 cytoplasmic puncta are distinct from persistent TiaR puncta

A prevailing hypothesis in the ALS field is that persistent stress granules seed TDP-43 inclusions leading to motor neuron death^9^. Notably, we observed colocalization of TDP-43 cytoplasmic puncta with TDP-43 phosphorylated at S409/410, but no obvious overlap with the autophagy protein p62 (***Fig. 4A, B***)^42^. We leveraged our recurrent stress paradigm to directly evaluate whether cytoplasmic puncta of TDP-43 and TiaR are related. In both NTg and TDP-43^M337V^ mice, TDP-43 puncta and TiaR puncta were observed in different neurons, with only 5-12% of neurons displaying both TDP-43 and TiaR puncta (***Fig. 4C, D*)**. Importantly, when TDP-43 puncta and TiaR puncta were detected in the same neuron, they were spatially distinct (***Fig. 4E, F***). Additionally, TDP-43 puncta did not overlap with the stress granule resident protein HuR (***Fig. 4E, F***). Our data indicate a clear distinction between cytoplasmic puncta that label for TDP-43 versus other RNA binding proteins using a recurrent heat stress paradigm *in vivo*.

**Figure 4.**
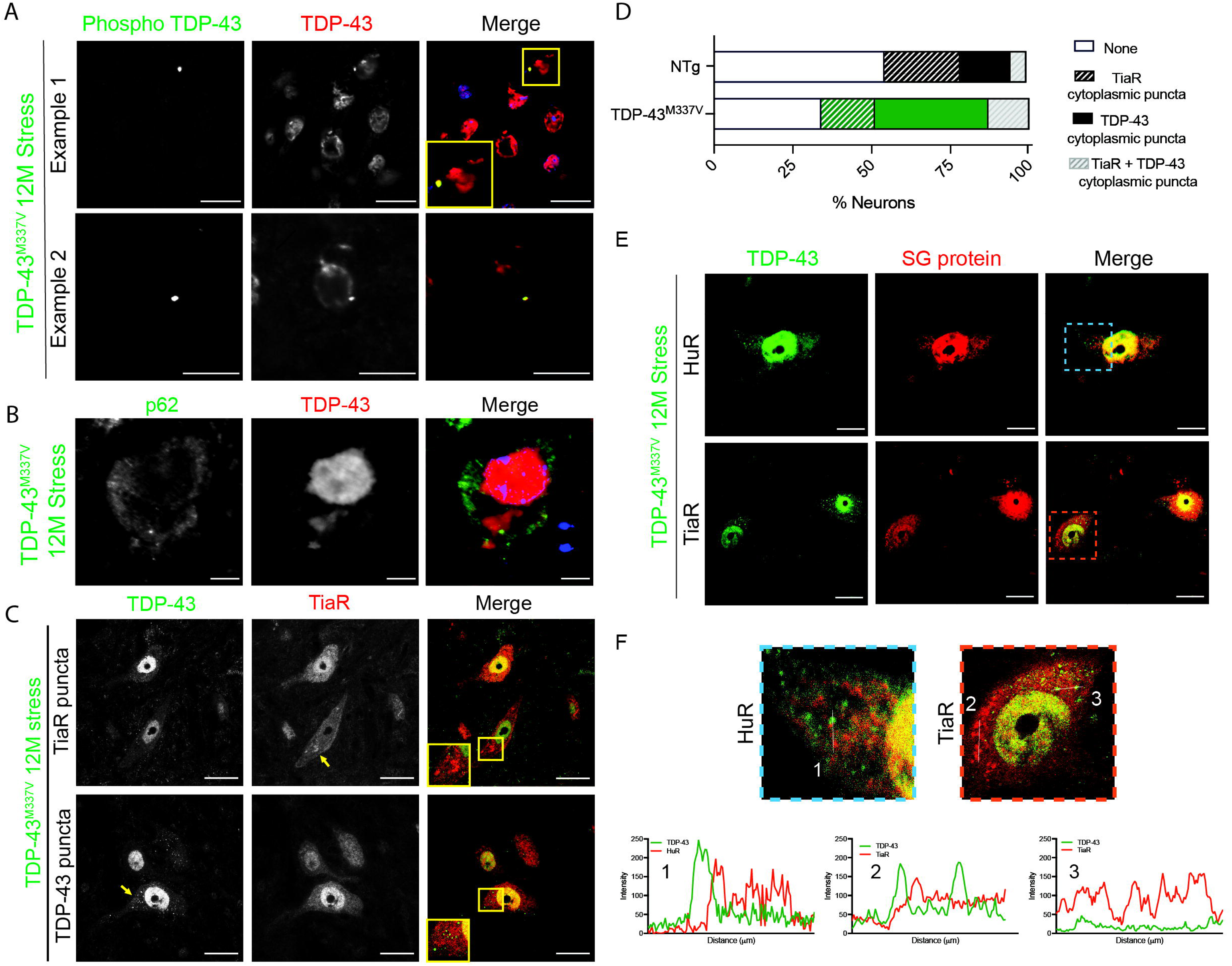
TDP-43 puncta resemble pathological inclusions and are independent of stress granule proteins. **(A)** Representative images showing overlap between TDP-43 inclusions with Phospho-TDP-43 (Ser 409/410) in 12M TDP-43^M337V^ mice after recurrent stress. Scale bar 20 µm (Example 1), 10 µm (Example 2). **(B)** Representative image of p62 and TDP-43 in 12M TDP-43^M337V^ mice after recurrent stress. Scale bar 5 µm. **(C)** Representative images with neurons after recurrent stress exhibiting either TDP-43 (green) or TiaR (red) puncta. Scale bar 10 µm. **(D)** Analysis of relationship between TDP-43 puncta formation and TiaR puncta formation. NTg (N=111); TDP-43^M337V^ (N=111). **(E)** Representative images with neurons containing stress granule protein puncta (TiaR, HuR) that do not overlap with TDP-43 puncta. **(F)** Line scans showing partitioning of green (TDP-43) and red (SG protein) signals in zoomed images.

### Selective alpha motor neuron death in TDP-43^M337V^ mice following recurrent stress

We next examined whether heat-induced cytoplasmic TDP-43 or TiaR puncta, and nuclear TDP-43 clearance, were associated with motor neuron loss. To distinguish α- and γ-motor neurons, spinal cord sections were co-labelled for ChAT and NeuN. As has been reported^43^, α-motor neurons express both ChAT and NeuN, whereas γ-motor neurons express ChAT, but not NeuN (***Fig. 5A, B*)**. Following recurrent hyperthermia in 4M and 12M mice, total motor neuron counts (ChAT⁺) appeared comparable in NTg mice, while there was a negligible trend for stressed TDP-43^M337V^ animals to have less motor neurons than sham-treated controls (***Fig. 5C, D***). However, closer inspection of α- and γ-motor neurons revealed a significant 23% reduction in α-motor neurons exclusively in 12M hyperthermia-treated TDP-43^M337V^ mice compared to sham controls with no significant change in γ-motor neuron numbers between conditions (***Fig. 5E, F***). There was no evidence of α- or γ-motor neuron loss in NTg mice at 4M or 12M (***Supplementary Fig. 5A, B***).

**Figure 5.**
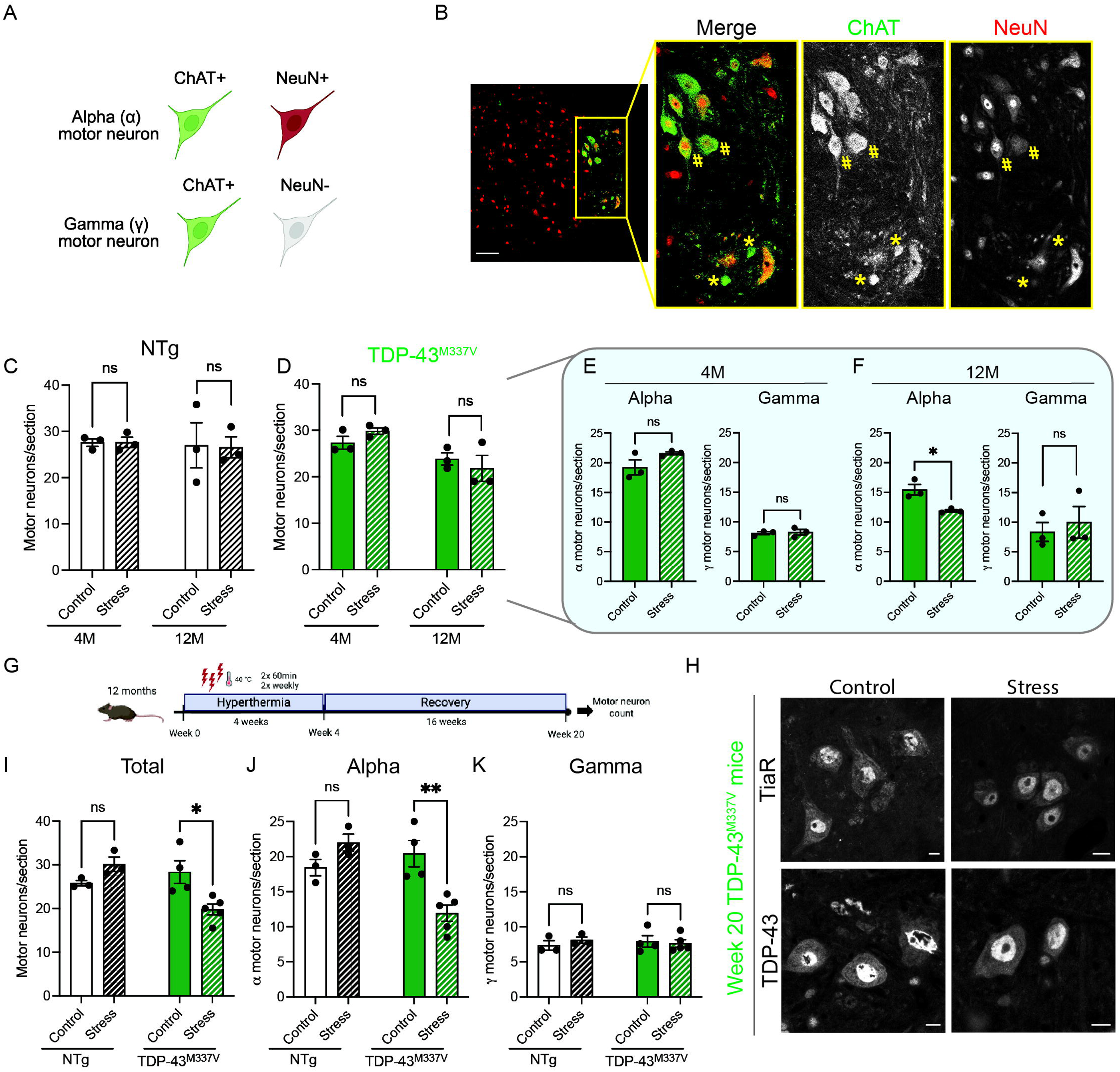
Selective, stress-induced, progressive motor neuron loss in TDP-43^M337V^ mice after hyperthermia. **(A)** Cartoon showing difference between alpha (α) and gamma (γ) motor neurons **(B)** Representative image of the ventral horn of a spinal cord to differentiate alpha and gamma motor neurons. Alpha (α) motor neurons are indicated with a hashtag (#) and gamma (γ) motor neurons are indicated with a star (*). Scale bar 50 µm. Total motor neuron counts in NTg **(C)** and TDP-43^M337V^ **(D)** mice at 4M and 12M following recurrent stress. **(E-F)** α and γ motor neuron counts in TDP-43^M337V^ mice at 4M and 12M. **(G)** Diagram of the stress and recovery paradigm used in 12M animals. **(H)** Representative picutres of TDP-43 and TiaR following a recovery period in 12M TDP-43^M337V^ mice. Scale bars 10 µm. Quantification of total **(I),** alpha **(J),** and gamma **(K)** motor neurons following a recovery period in NTg and TDP-43^M337V^ mice. For all graphs, each point represents a single mouse. One-way ANOVA analysis was used to determine statistical significance.

To determine whether these effects persisted after stress recovery, a separate cohort of 12M TDP-43^M337V^ and NTg mice were subjected to recurrent hyperthermia for 4 weeks followed by a 16 week recovery period (***Fig. 5G***). In this recovery paradigm, no persistent stress granules were observed at 20 weeks as assessed by polyA mRNA signal (***Supplementary Fig. 5C***). Additionally, neither cytoplasmic TiaR puncta, nor cytoplasmic TDP-43 puncta and nuclear clearance were detected in mice of either genotype (***Fig. 5H, Supplementary Fig. 5D***). At week 20, total motor neuron counts were reduced by 30% in TDP-43^M337V^ mice (***Fig. 5I***), driven entirely by a selective 42% loss of α-motor neurons (***Fig. 5J***), while γ-motor neurons remained unaffected (***Fig. 5K***). In contrast, motor neuron counts were unaffected by recurrent heat stress in NTg littermates, neither immediately after heat exposure (***Fig. 5B, Supplementary Fig. 5***) or following 16 weeks of recovery (***Fig. 5I***).

Taken together, these results indicate that recurrent hyperthermia induces cytoplasmic TiaR puncta that are reversible and not associated with motor neuron death, as they are detected in both NTg and TDP-43^M337V^ mice and resolve during recovery. However, in TDP-43^M337V^ mice, we note that a subset of neurons exhibiting cytoplasmic TDP-43 puncta (40%), which are not detectable after recovery, parallel a subset of neurons that are lost (44%). We infer that the neurons which were lost are the same neurons that displayed TDP-43 cytoplasmic puncta. Thus, recurrent hyperthermia in genetically predisposed mice reproduces key features of ALS pathology, including nuclear TDP-43 clearance, cytoplasmic TDP-43 puncta formation, and selective vulnerability of α-motor neurons.

## Discussion

The hypothesis that stress granules act as crucibles for TDP-43 inclusions, which trigger motor neuron death in ALS, is largely based on *in vitro* studies that rely on overexpression of aggregation-prone proteins^10,44,45^. Here, we employed a recurrent hyperthermia model and a mouse model expressing near-physiological levels of human mutant TDP-43 protein to probe this relationship *in vivo*. The homozygous TDP-43^M337V^ mouse model used in this study exhibits a 1.1x-fold expression of human TDP-43 compared to endogenous mouse Tdp-43. This model permits interrogation of the consequence of this disease-associated mutation, avoiding the pitfalls of gross overexpression. Using 60 mins hyperthermia exposure in awake, restrained mice, we observed stress granule formation and dissolution in lumbar motor neurons, as assessed by polyA mRNA and canonical stress granule protein markers. Extending the 60 mins stress to a recurrent paradigm allowed us to observe that while polyA mRNA-positive granules did not persist after 16 hyperthermia exposures delivered over the course of one month, puncta labelling for stress granule-resident proteins TiaR, G3BP2, and HuR did. Interestingly, young (4M) TDP-43^M337V^ mice displayed a similar frequency of TiaR puncta as NTg animals, despite their known stress granule assembly defect^16^. The concept that stress granule proteins can assemble into RNA-deficient aggregates is known^46^. For example, poliovirus-infected cells form Tia1 and TiaR positive aggregates that are distinct from stress granules, since they lack polyadenylated mRNA^46^. Notably, poliovirus induces cell death in part by suppressing the stress granule response^47^. This phenomenon loosely parallels our observations in TDP-43^M337V^ mice, which fail to form stress granules yet still exhibit TiaR-positive cytoplasmic puncta. Similar to stress granules, phase-separated TDP-43 is thought to be stabilized by RNA interactions that limit its cytoplasmic aggregation^48–50^. In our recurrent stress paradigm, the absence of polyadenylated mRNA puncta, together with positive staining using an aptamer that selectively recognizes RNA-deficient TDP-43^41^, suggests that the TDP-43 puncta formed under these conditions more closely resemble aggregates than dynamic phase-separated assemblies. Importantly, hyperthermia-induced persistent TiaR puncta do not correlate with motor neuron loss, indicating that the persistence of stress granule protein puncta is not inherently pathological as they are also observed in NTg animals that have no motor neuron loss.

In aged (12M) TDP-43^M337V^ mice exposed to recurrent hyperthermia, we observed both nuclear clearance of TDP-43 and cytoplasmic puncta formation, but strikingly, these events were mutually exclusive within individual neurons and independent of the formation of polyA mRNA-positive stress granules. Additionally, TDP-43 puncta were observed to be spatially distinct of TiaR or HuR puncta, which does not support a recent report, using overexpression of a cytoplasmic-restricted TDP-43 variant (TDP-43^ΔNLS^), claiming that stress granules scaffold TDP-43 phase separation^8,51^. We appreciate that our observations capture static endpoints after one month of stress, and thus we cannot exclude sequential or overlapping events during the recurrent hyperthermia paradigm. However, our data are consistent with other work that demonstrates that demixing of TDP-43 puncta from stress granules is associated with cell death and suggests that this separation is a key driver of the cell death phenotype^52^.

Similar to many TDP-43 mouse models that have near physiological levels of TDP-43, TDP-43^M337V^ mice exhibit no motor neuron loss, as assessed up to 24M^29^. However, after recurrent hyperthermia, these mice developed selective alpha motor neuron loss, which worsened during a recovery period. Intriguingly, motor neuron loss was inversely correlated with TDP-43 puncta frequency. At 4M and 12M, translocation and puncta formation were observed without significant neuron loss but after a recovery period, motor neuron death was evident while TDP-43 localization was primarily nuclear in surviving neurons. This suggests that in this model, either TDP-43 nuclear clearance or puncta formation can trigger neurodegeneration, though we cannot yet distinguish their relative toxicity.

### Reproducing the genetics x time x environment model of ALS

To integrate our findings within a broader framework, we refer to the model proposed by Al-Chalabi and Hardiman ^17^, which conceptualizes ALS as arising from the intersection of genetic susceptibility, aging, and environmental exposures. In our study, we modeled these components using aged TDP-43^M337V^ mice subjected to recurrent hypothermia exposures (***Fig. 6***). Unlike TDP-43 overexpression models with severe phenotypes^30,53^, TDP-43 models with physiological expression show only mild FTD-like, but not motor, deficits^19,20^. Our findings here support the idea that these models require a “second hit” to unmask neurodegenerative processes. In our system, aging provides a third factor, leading to a “three-hit” model where genetic predisposition, environmental stress, and aging (time) converge to elicit ALS-like pathology, similar to human ALS^17^. Consistent with prior *in vitro* observations^54^, our data also suggest that nuclear TDP-43 clearance precedes cytoplasmic puncta formation in the context of ALS burden. Finally, it is important to note that this “three-hit” model that reproduces major features of ALS pathophysiology emerges in the context of defective stress granule assembly. We propose that this recurrent hyperthermia model can be leveraged to study early TDP-43-mediated disease mechanisms and to evaluate preclinical therapeutics targeting TDP-43 mislocalization or loss of function.

**Figure 6.**
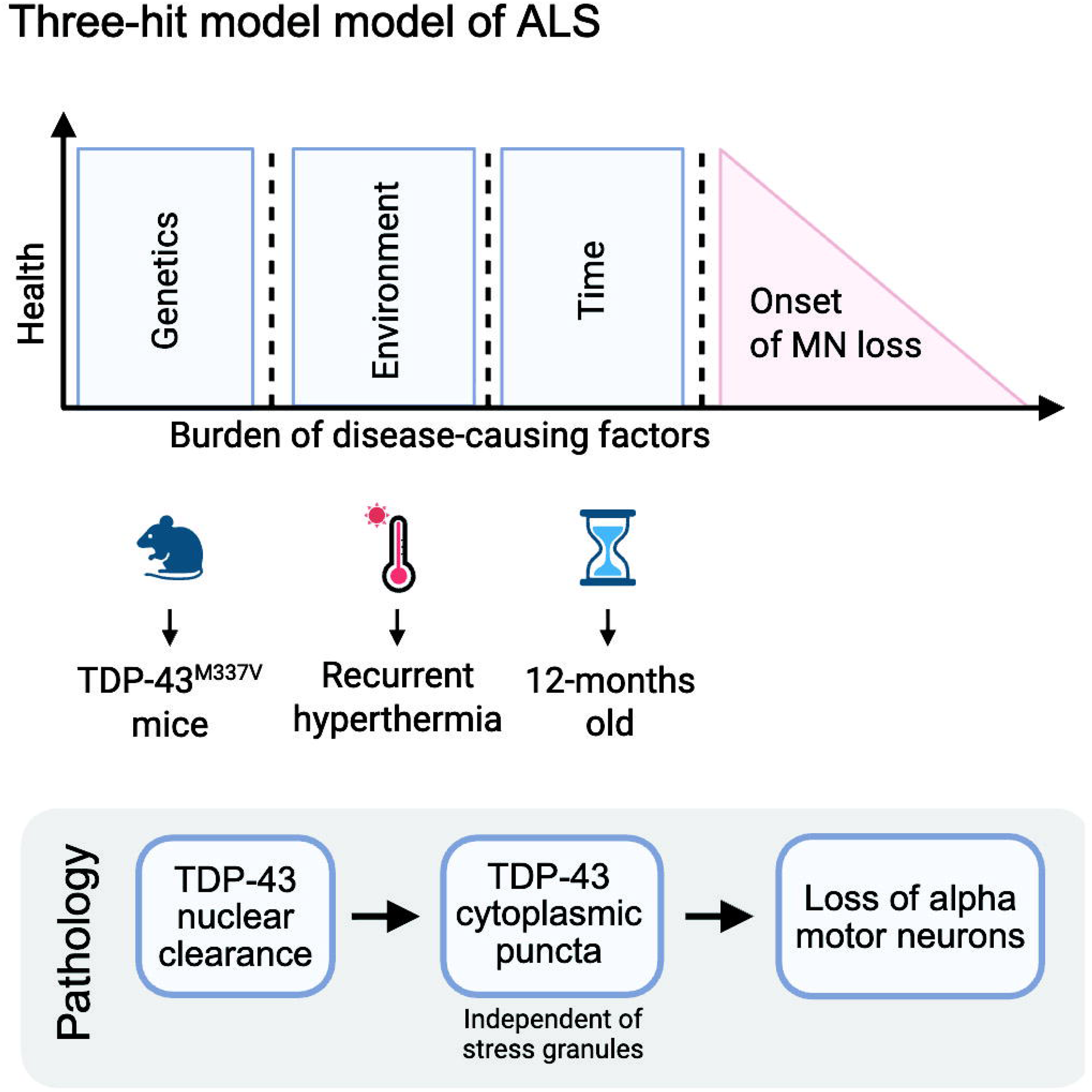
A model recapitulating the genes, time, environment hypothesis for ALS. Model showing the genes, time, environment hypothesis in the context of this study using recurrent stress on 12M TDP-43^M337V^ mice. The sequence of TDP-43 pathology and motor neuron loss is also indicated.

## Conclusion

In this study, we leveraged a recurrent hyperthermia paradigm to demonstrate that persistent, stress-induced TiaR puncta are not toxic to spinal motor neurons in aged mice *in vivo* and do not spatially or temporally correlate with TDP-43 puncta following repeated stress. Consistent with this dissociation, we show that in the TDP-43^M337V^ mouse model where stress granule assembly is impaired, stress exposure nonetheless induces TDP-43 nuclear clearance and cytoplasmic puncta accumulation. This TDP-43 disease signature is accompanied by a progressive and selective loss of spinal α-motor neurons during cycles of stress and recovery. Importantly, the presence of motor neuron degeneration demonstrates that TDP-43 mislocalization, rather than the persistence of TiaR puncta, correlates with selective α-motor neuron vulnerability in the spinal cord. This is evidence that TDP-43 puncta arise independent of canonical stress granule proteins in a mouse model expressing physiologically relevant levels of TDP-43 and directly challenges the idea that stress granule inhibition is a therapeutic avenue. Lastly, the presented data indicate that the convergence of genetic susceptibility, environmental stress, and time is sufficient to drive both molecular and cellular hallmarks of neurodegenerative disease.

## Declarations

### Ethics

The use of animals and all procedures were performed in according to guidelines of the Canadian Council on Animal Care and were approved by the CRCHUM animal care committee (CIPA).

### Availability of data and materials

The data that support the findings of this study are available from the corresponding author, upon reasonable request.

## Author contributions

AD and CVV designed and conceptualized the study. AF, MC, HS, AA, CF, MT, MG, and ML contributed to the experimental work. SP managed the animal colonies and provided oversight for animal protocols. JG provided oversight to aptamer work. AD and CVV wrote the manuscript. All authors reviewed and approved the manuscript.

## Acknowledgements

We thank all CVV lab members for helpful insights and discussion. We also thank Kevin Talbot for providing the TDP-43^M337V^ mice and Aurélie Cleret-Buhot from the CRCHUM imaging platform for technical support.

## Supplemental figure legends

**Supplementary Fig. 1. TDP-43^M337V^ mice have a stress granule defect after 60 mins of hyperthermia.** Representative images of polyA mRNA at T=0, T=60, and T=75 in 4M TDP-43^M337V^ mice (N=3 per time point). 60 min stress was done at 40°C exactly like in NTg mice. Scale bars 10 µm.

**Supplementary Fig. 2. Temperature and weight during recurrent hyperthermia.** Average temperature for each mouse during a 60 mins stress; weight loss over 4 weeks of recurrent hyperthermia for mice starting the paradigm at **(A)** 4M and **(B)** 12M.

**Supplementary Fig. 3. G3BP2 and HuR puncta persist after recurrent hyperthermia.** Representative images of overlap between HuR and G3BP2 puncta in neurons from 4M NTg mice after stress. Scale bar 10 µm.

**Supplementary Fig. 4. Presence of TDP-43 cytoplasmic puncta in 12M TDP-43^M337V^ mice after recurrent stress. (A)** Additional images showing TDP-43 cytoplasmic inclusions with the Proteintech and R&D Biosystems antibodies in 12M TDP-43^M337V^ mice that underwent recurrent hyperthermia. Scale bars 10 µm. **(B)** Quantification of neurons with TDP-43 puncta at 12M using Proteintech antibody. NTg: Control 12M (N=3), Stress 12M (N=4), TDP-43^M337V^: Control 12M (N=4), Stress 12M (N=4). **(C)** Analysis of relationship between TDP-43 nuclear clearance and puncta formation with Proteintech antibody. NTg (N=101); TDP-43^M337V^ (N=100). **(D)** RT-PCR depicting cryptic *Adnp2*, cryptic *Synjbp2*, cryptic *Usp15*, and *Sort1* exon17B in NTg and TDP-43^M337V^ mice following recurrent stress.

**Supplementary Fig. 5. Alpha and gamma motor neurons are unchanged with recurrent hyperthermia in non-transgenic mice.** Quantification of α-motor neurons and γ-motor neurons at 4M **(A)** and 12M **(B)** in NTg mice following recurrent hyperthermia. For all graphs, each point represents a single mouse. One-way ANOVA analysis was used to determine statistical significance. **(C)** Representative polyA mRNA FISH images for NTg and TDP-43^M337V^ mice following a recovery period after the 12M recurrent hyperthermia treatment. Scale bars 10 µm. **(D)** Representative images of TiaR and TDP-43 in 12M NTg mice following a recovery period. Scale bars 10 µm.

