## Supplementary figures and images for "TDP-43 pathology is linked to motor neuron loss and is independent of stress granules *in vivo*"

### Supplemental Figures

FIGURE S1

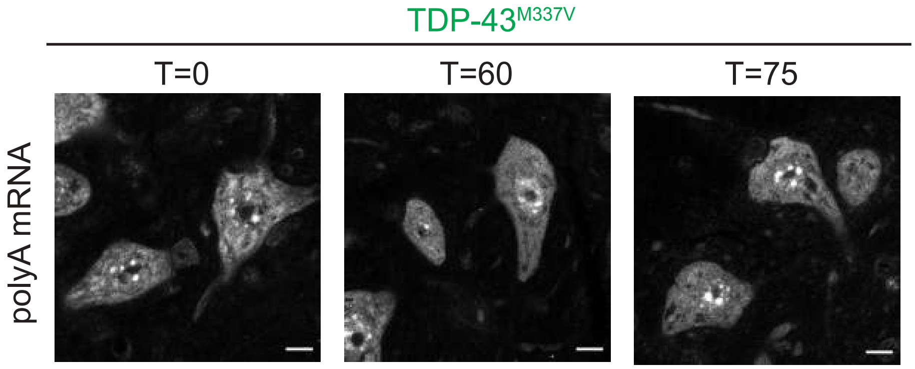

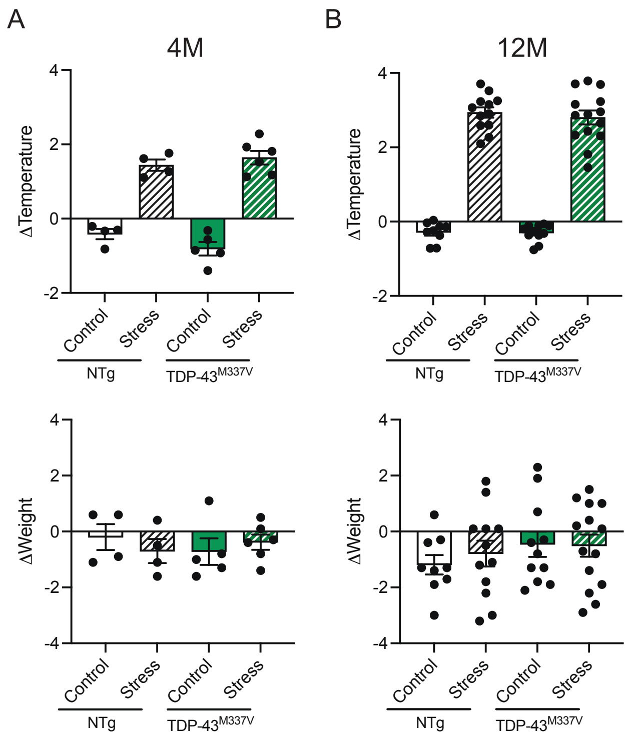

FIGURE S3

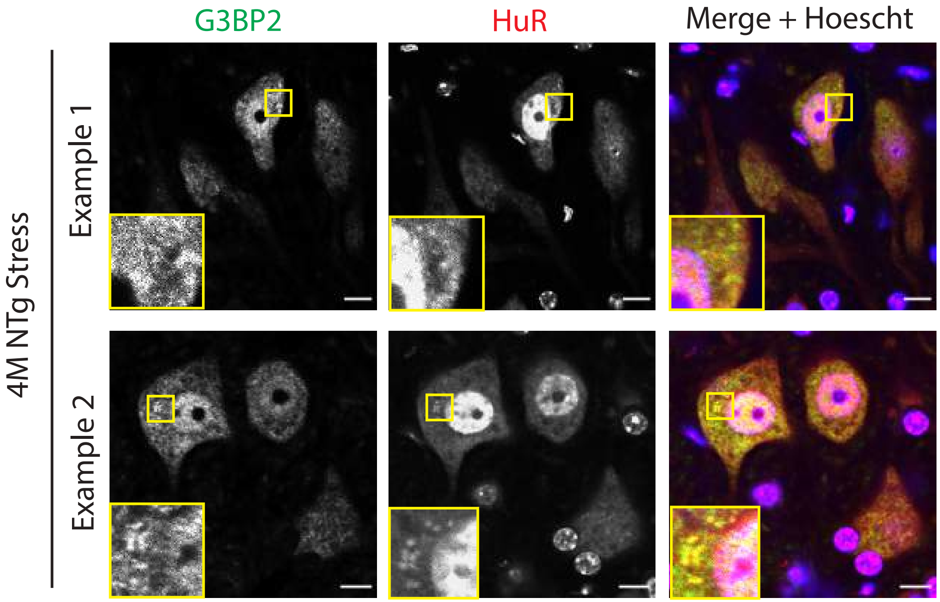

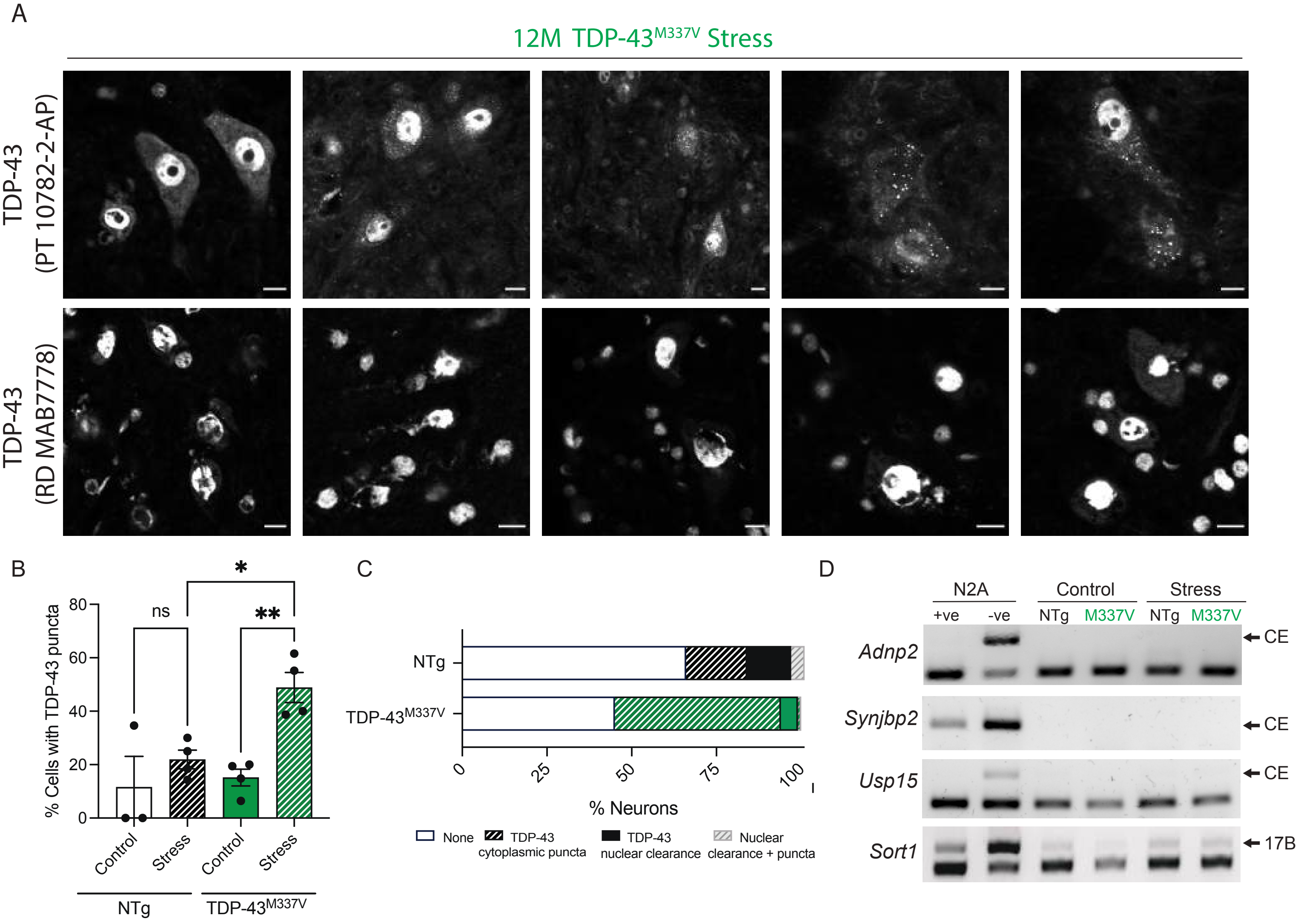

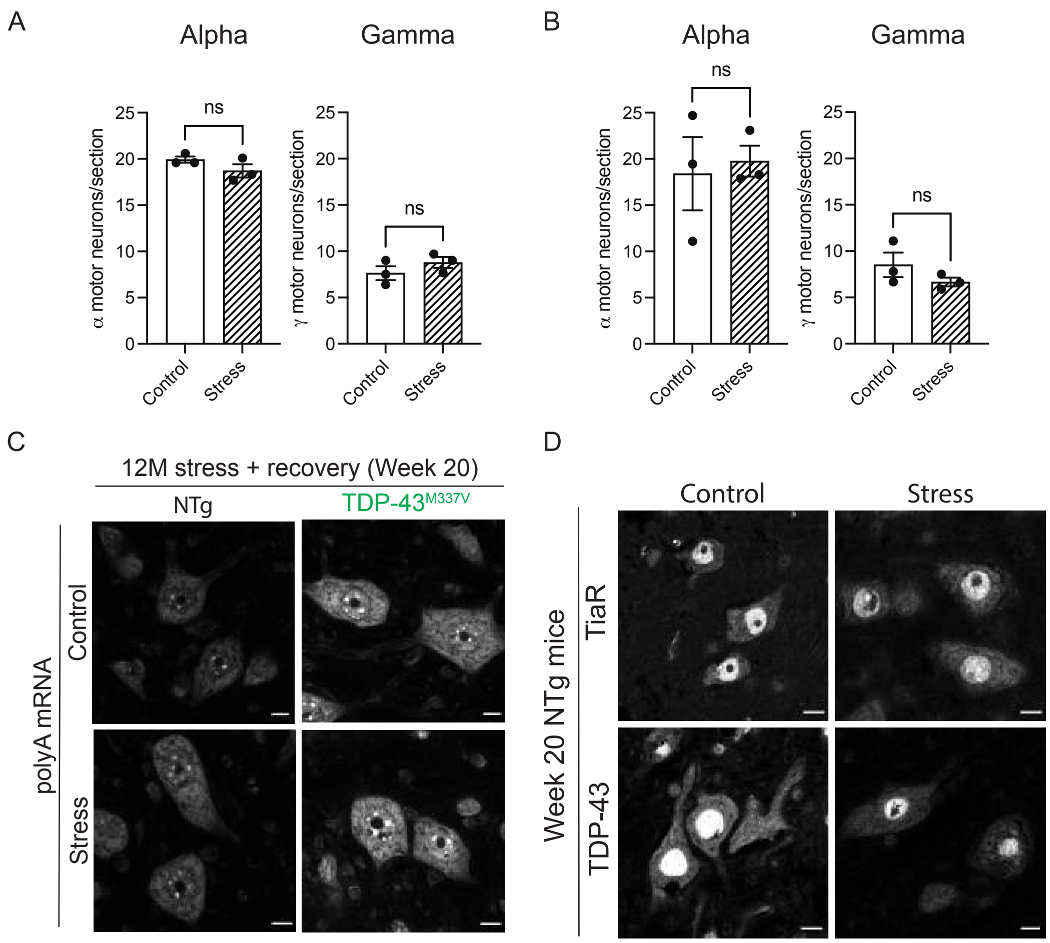
